# The C-terminal domain of *Clostridioides difficile* TcdC is exposed on the bacterial cell surface

**DOI:** 10.1101/872283

**Authors:** Ana M. Oliveira Paiva, Leen de Jong, Annemieke H. Friggen, Wiep Klaas Smits, Jeroen Corver

## Abstract

*Clostridioides difficile* is an anaerobic gram-positive bacterium that can can produce the large clostridial toxins, Toxin A and Toxin B, encoded within the pathogenicity locus (PaLoc). The PaLoc also encodes the sigma factor TcdR, that positively regulates toxin gene expression, and TcdC, a putative negative regulator of toxin expression. TcdC is proposed to be an anti-sigma factor, however, several studies failed to show an association between *tcdC* genotype and toxin production. Consequently, TcdC function is not yet fully understood. Previous studies have characterized TcdC as a membrane-associated protein with the ability to bind G-quadruplex structures. The binding to the DNA secondary structures is mediated through the OB-fold domain present at the C-terminus of the protein. This domain was previously also proposed to be responsible for the inhibitory effect on toxin gene expression, implicating a cytoplasmic localization of the C-terminal OB-fold.

In this study we aimed to obtain topological information on the C-terminus of TcdC. Using Scanning Cysteine Accessibility Mutagenesis and a HiBiT-based system, we demonstrate that the C-terminus of TcdC is located extracellularly. The extracellular location of TcdC is not compatible with direct binding of the OB-fold domain to intracellular nucleic acid or protein targets, and suggests a mechanism of action that is different from characterized anti-sigma factors.

**Importance:** Transcription of the *C. difficile* large clostrididial toxins (TcdA and TcdB) is directed by the sigma factor TcdR. TcdC has been implicated as a negative regulator, possible acting as an anti-sigma factor.

Activity of TcdC has been mapped to its C-terminal OB fold domain. TcdC is anchored in the bacterial membrane, through its hydrophobic N-terminus and acting as an anti-sigma factor would require cytoplasmic localization of the C-terminal domain.

Remarkably, topology predictions for TcdC suggest the N-terminus to be membrane localized and the C-terminal domain to be located extracellularly. Using independent assays, we show that the C-terminus of TcdC indeed is located in the extracellular environment, which is incompatible with its proposed role as anti-sigma factor in toxin regulation.

## Introduction

*Clostridioides difficile (Clostridium difficile*) (1) is an opportunistic pathogen that can cause disease in individuals with dysbiosis of the gut microbiota (2). *Clostridium difficile* infection (CDI) incidence has increased worldwide and leads to a broad spectrum of symptoms, from mild diarrhea to toxin megacolon, and even death (3).

Several factors contribute to the progression and the severity of CDI (2, 4). *C. difficile* is a Gram-positive anaerobic bacterium that has the ability to form spores, which allows for dissemination and colonization (2). The main virulence factors are the large clostridial toxins that induce damage to the epithelial cells and lead to an inflammatory response that underlies the symptoms of CDI (2, 3, 5).

*C. difficile* strains have been found to produce up to three toxins: Toxin A (TcdA), Toxin B (TcdB) and binary toxin (CDT) (5, 6). Toxins A and B are encoded by genes *tcdA* and *tcdB*, respectively, located on a 19.6 kb chromosomal region termed pathogenicity locus (PaLoc) (7). TcdA and TcdB are glucosyltransferases and once translocated to the cytosol of the intestinal epithelial cells, start a cascade of events that can eventually lead to cell death (2, 5). CDT, encoded by the *cdtA* and *ctdB* genes, is an ADP-ribosylating toxin that acts on the actin cytoskeleton (8).

The PaLoc contains at least 3 additional genes that appear to be involved in the regulation of the expression or function of the large clostridial toxins: *tcdE, tcdR* and *tcdC* (5, 6). TcdE is a putative holin-like protein, thought to be involved in toxin secretion, however its exact role is still unclear (9). TcdR is an RNA polymerase sigma factor that acts as the positive transcriptional regulator of *tcdA, tcdB* and *tcdE* and also positively regulates its own expression. A direct interaction between TcdR and RNA polymerase allows the recognition of the target promoters and activates expression (5, 10). Expression of *tcdR,* and consequently *tcdA* and *tcdB,* is influenced by different stimuli, such as temperature, nutrient availability and medium composition (11-14).

Analysis of gene transcription by quantitative PCR has shown that while the expression of *tcdA, tcdB, tcdE* and *tcdR* is low during exponential phase, these strongly increase upon entering stationary phase. In contrast, *tcdC* was found to be highly expressed during exponential phase but to decrease in stationary phase (15). Similar profiles were shown at the protein level, where levels of TcdC were higher in the exponential growth phase (16). Together, these data suggested that TcdC might act as a negative regulator of toxin transcription. However, several other studies did not find a decrease in *tcdC* transcription in stationary phase and but rather showed a constant expression level during the stationary growth phase (14, 17, 18).

Likewise, the association between toxin expression and *tcdC* gene variants is subject of debate. Increased virulence in epidemic strains was thought to be caused by deletions and frameshift mutations in *tcdC,* leading to a severely truncated non-functional protein and presumably higher toxin titers as a consequence (19, 20). In support of this, it was shown that introduction of a plasmid-based copy of the wild type *tcdC* gene in strain M7404 (PCR ribotype 027, carrying a truncated *tcdC*) resulted in decreased virulence in hamsters (20). However, mutations in the *tcdC* gene of clinical isolates did not predict the activity of toxins A and B (18, 21). Moreover, several studies failed to observe a relation between toxin gene expression and *tcdC* genotype. Restoration of chromosomal *tcdC* of outbreak strain R20291 (PCR ribotype 027) to wild type did not result in altered toxin expression (22) and toxin expression in *C. difficile* 630Δ*erm* and an isogenic *tcdC* clostron mutant showed no significant differences observed in toxin levels (17).

Previous studies have characterized the domain structure of TcdC (16, 23). TcdC is a 26 kDa dimeric protein that contains an N-terminal transmembrane region (residues 30-50), that allows its anchoring to the cell membrane, a coiled-coil dimerization domain and a C-terminal functional domain (10, 23). Using surface plasmon resonance experiments, purified full length TcdC was shown to interact with *E.coli* core RNA polymerase and prevented the formation of the active holoenzyme TcdR-RNA polymerase (10). Overexpression of *C. difficile* TcdC in the heterologous host *Clostridium perfringens* results in repression of TcdR-driven transcription from the *tcdA* promoter, and the C-terminal domain of TcdC was sufficient for this activity (10). However, it is not clear if TcdR and TcdC are in close proximity inside the bacterial cell.

Due to lack of structural characterization of TcdC homologues, computational analysis was used to build a structural model of the C-terminal domain of TcdC. This modelling suggested the domain adopts a dimeric, ssDNA-binding OB-fold (Oligonucleotide/Oligosaccharide Binding fold) (23). TcdC is capable of binding to ssDNA G-quadruplexes *in vitro*, but considering the paucity of these structures in the genome sequence of *C. difficile*, G-quadruplexes might mimic an alternative TcdC binding partner (23).

It is clear that further studies are required to understand TcdC binding partners and their function in transcriptional repression. The prevailing model is that TcdC functions as an anti-sigma factor, whose activity depends on cytosolic localization of the C-terminal OB-fold domain. However, the topological information of the C-terminal domain has not been demonstrated to date.

In this study, we aimed to determine whether the C-terminal domain of TcdC is cytosolic or surface exposed. Using different topology prediction methods, Substituted Cysteine Accessibility Method (SCAM) and a codon-optimized version of the HiBiT Extracellular Detection System (HiBiT^opt^), we find that the C-terminal domain of TcdC is located extracellularly.

## Results

### *In-silico* prediction of TcdC topology suggests an extracellular location of the C-terminal domain

To analyze the topology of *C. difficile* TcdC (CD0664 from *C. difficile* 630) we first analyzed the protein sequence (Uniprot ID: Q189K7) using three different prediction algorithms: TMHMM 2.0 (http://www.cbs.dtu.dk/services/TMHMM-2.0) (24), TOPCONS 2.0 (http://topcons.cbr.su.se/) (25) and SignalP 5.0 (http://www.cbs.dtu.dk/services/SignalP/) (26).

TMHMM 2.0 (24) predicts a transmembrane helix of around 16 residues (residues 31-46) with moderate probability. Residues 1-13 are predicted to be inside of the cell (0.63 probability) whereas the C-terminal region (coiled coil and OB-fold domains) is predicted to be outside of the cell (probability >0.8) (Fig 1A). The consensus of TMHMM 2.0 1-best algorithm predicts TcdC to be extracellular (Fig. 1A, pink bar). TOPCONS 2.0 (25), which identifies regions with a low free energy difference (ΔG), similarly suggests the presence of an N-terminal transmembrane helix. The TOPCONS 2.0 consensus prediction is an intracellular N-terminal domain (residues 1 – 26), a transmembrane helix (residues 27 – 46) and an extracellular C-terminal region that encompasses the dimerization and OB-fold domains (residues 47 – 232).

**Figure 1.**
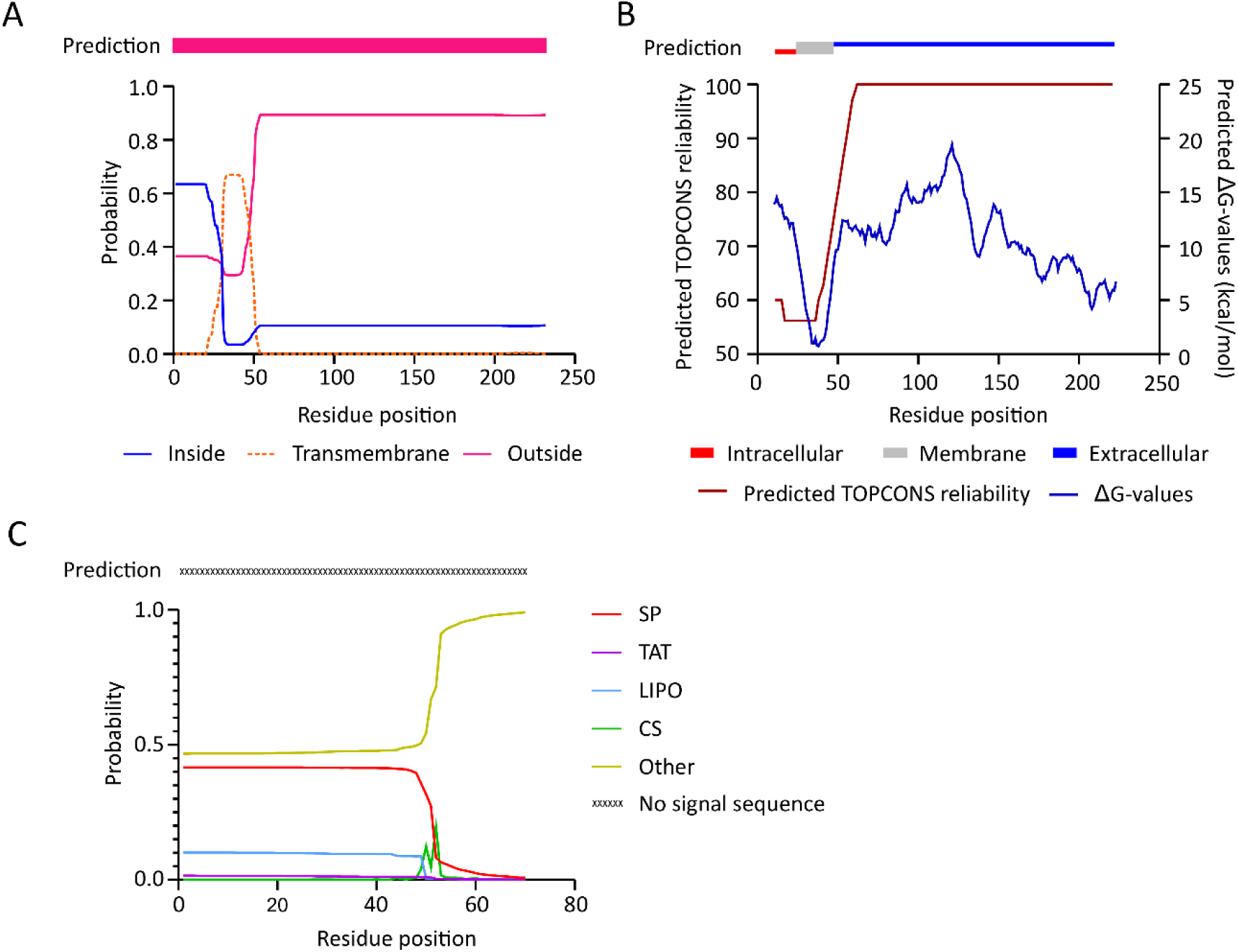
Prediction of a transmembrane helix in TcdC. **A)** Output from the prediction by TMHMM 2.0 software (24) through 1-best algorithm (pink bar) and probability plot: inside the cell (blue line), transmembrane region (orange dotted line) and outside the cell (pink line). **B)** Prediction by TOPCONS software (25), with consensus in residues 1-26 inside the cell (red box), a transmembrane helix (residues 26-46, grey box) and residues 47-232 on the outside of the cell (blue line). TOPCONS reliability score (brown line) and predicted ΔG-values for each residue (blue line) as shown. **C)** Output from the SignalP 5.0 web-server for the TcdC amino acid sequence. No signal sequence was detected (X). Probabilities of signal peptides presence from the systems Sec (SP, red line), Tat (purple line), and lipoprotein (LIPO, blue line) are shown. Predicted cleavage site score (CS, green line) and no signal sequence probability is depicted (OTHER, light green line).

We also investigated the presence of a potential signal peptide in the TcdC amino acid sequence through SignalP 5.0 (26). However, no known signal peptide was identified, suggesting TcdC remains tethered to the membrane (Fig. 1C).

Though the reliability of the predictions is relatively low, both TMHMM and TOPCONS support the presence of the transmembrane helix (Fig.1), consistent with previous observations (10, 16). Strikingly, both methods suggest that the TcdC C-terminus is located outside of the cell. As this would be incompatible with a role for the OB-fold domain in sequestering TcdR or repression of TcdR-mediated transcriptional activation, we set out to obtain topological information on the OB-fold domain in *C. difficile*.

### TcdC C-terminus is accessible for extracellular cysteine labelling

We performed the Substituted Cysteine Accessibility Method (SCAM) to evaluate the location of the TcdC C-terminus. SCAM subjects cysteine residues present in the protein of interest to chemical modification with the thiol-specific probe N-(*3*-maleimidylpropionyl) biocytin (MPB), that has a low membrane permeability. The probe forms a stable, non-hydrolysable bond with the thiol-group of a cysteine residue, resulting in the biotinylation of the protein. At low concentrations MPB exclusively labels extracytoplasmic (surface exposed) cysteines, providing topological information about the labelled protein (27).

A typical SCAM experiment relies on immunoprecipitation of protein (using antibody specific for the protein of interest), detection of immunoprecipitated protein (using a second antibody, directed at a tag on the protein of interest), and verification of labelling with the MPB (using anti-biotin antibodies). We introduced a C-terminally 3xmyc-tagged TcdC construct (TcdC-3xmyc), under the control of the inducible promoter P_tet_ (28), that can be precipitated with anti-TcdC (α-TcdC) antibody and detected using anti-myc antibodies (α-myc). We affinity purified a previously generated TcdC antibody (17) and verified its specificity on *C. difficile* lysates by immunoblotting. The TcdC-3xmyc construct was induced and samples before and after induction were analyzed. Only in the induced samples a band migrating at the approximate molecular weight of TcdC was observed (38 kDa, Fig. 2A), suggesting that native levels of TcdC under non-inducing conditions are below our limit of detection in this assay. Though the predicted molecular weight of the TcdC protein is 23 kDa, a higher molecular weight on SDS-PAGE has been observed before of approximately 37 kDa (10, 23). Several bands were detected with a lower molecular weight than expected (Fig. 2A). The fact that these are only present when TcdC-3xmyc is induced (Fig. 2B) suggests possible alternative forms of the protein. Nevertheless, the apparent specificity of the anti-TcdC antibody allowed further SCAM analysis.

**Figure 2.**
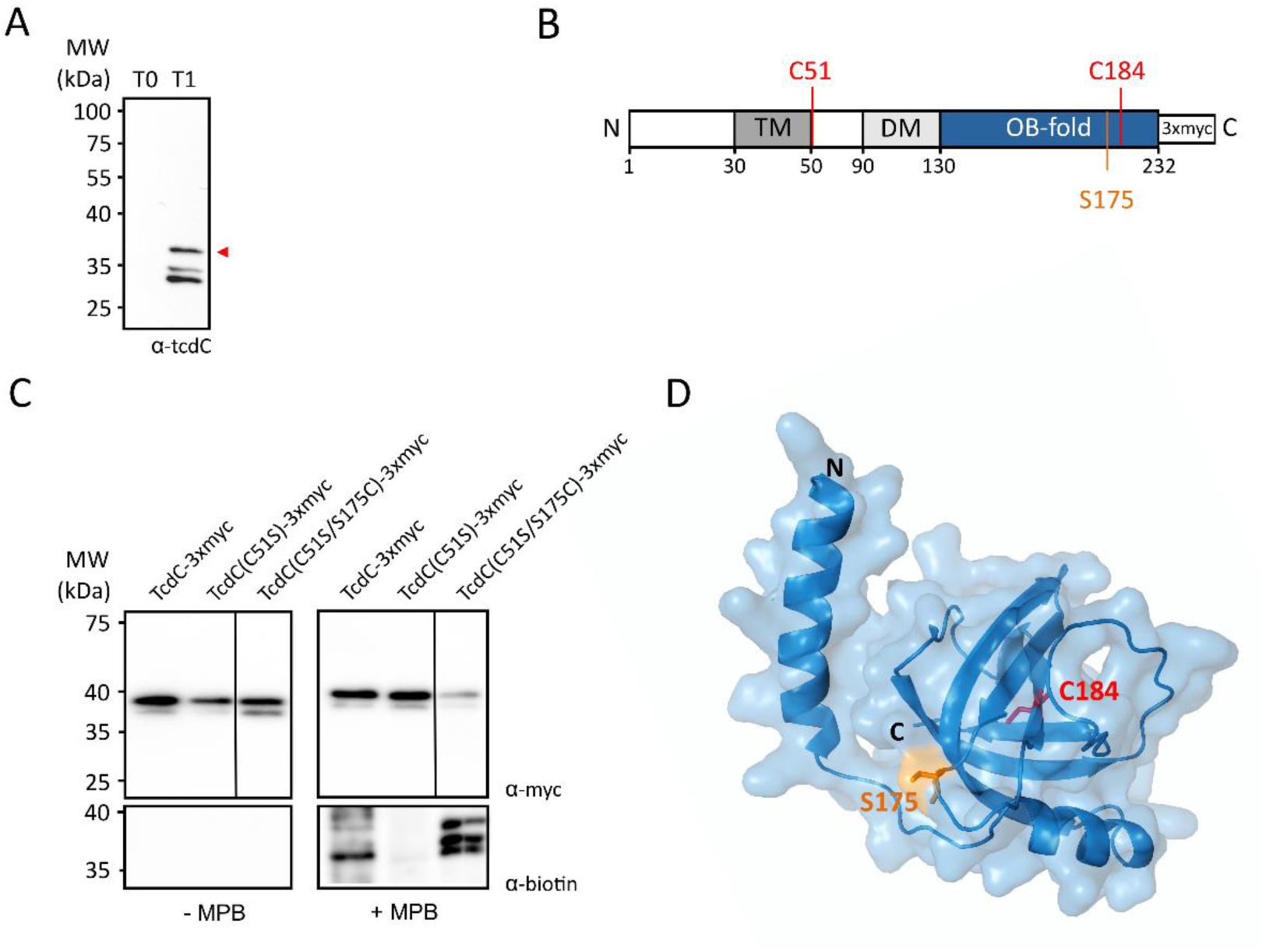
Mapping the location of the TcdC topology-terminus using SCAM. **A)** Western-blot analysis of α-TcdC antibody specificity in *C. difficile* lysates harbouring pLDJ1 (P_tet_-*tcdC-3xmyc*), before (T0) and after induction with 200ng/ml anhydrotetracycline for 2 hours (T2). Full-length TcdC is indicated with a red arrow. **B)** Schematic representation of the 3xmyc-tagged TcdC construct used for the SCAM analysis. The different domains of TcdC are represented: transmembrane domain (TM, grey box), the dimerization domain (DM, light grey box) and the OB-fold (blue box). The 3xmyc-tag is represented as a white box, the residues used in the SCAM analysis are represented in red (cysteines) and orange (serine). **C)** SCAM analysis of the different TcdC_3xmyc_ constructs. The strains harbouring the different myc-tagged TcdC constructs (38 kDa) were induced for 2 hours. Samples were collected and treated with 1 mM MPB (+, right two panels) or with no MPB (-, left two panels). Samples were immunoprecipitated and immunoblotted with α-myc for TcdC-3xmyc protein detection (upper panels), and α-biotin for detecting biotinylated proteins (bottom panels). **D)** Mapping of S175 and C184 residues on TcdC structural model. Structural model of the TcdC C-terminal previously determined by I-TASSER (residues 86 to 232; UniProt ID: Q189K7) (23) and structural visualization through PyMOL Molecular Graphics System, Version 1.76.6 Schrödinger, LLC. Both N- and C-terminus are marked in the figure. The C-terminal residues S175 (orange stick) and C184 (red stick) are indicated. S175 is exposed on the protein surface and C184 is buried inside the C-terminal OB-fold.

TcdC contains 2 endogenous cysteines; one at position 51, right after the predicted transmembrane domain (residues 30 to 50) and another one at position 184, located in the predicted OB-fold (Fig. 2B). To evaluate the cysteine labelling on the native protein, we assessed biotinylation of TcdC-3xmyc (Fig. 1C). The signal in the anti-biotin (α-biotin) Western blot suggests that one, or both, of the native cysteine residues in TcdC-3xmyc is accessible for labelling by MPB. To delineate which of the two cysteine residues (C51 and C184) contributes to the signal, we constructed a TcdC mutant in which one cysteine residue is changed into a serine residue. Mutation of C51 (TcdC(C51S)-3xmyc) resulted in loss of biotinylation in the presence of MPB (Fig. 1C), despite the fact that C184 was still present, suggesting that it is C51 that is biotinylated in TcdC-3xmyc. The lack of signal was not due to poor expression of the protein, as the signal in the anti-myc Western blot is comparable to the TcdC-3xmyc protein.

As both native cysteines were predicted to be outside of the cell (Fig. 1, Fig. 2B), we expected biotinylation of C184 in the C51S mutant. Careful investigation of the predicted OB-fold structure (Fig. 2D) suggested that C184 was buried in the OB fold, inaccessible for biotinylation. To determine if a failure to biotinylate is not a general feature of the OB-fold domain, we then introduced a cysteine in TcdC(C51S)-3xmyc at position 175 (TcdC(C51S/S175C)-3xmyc), located in a predicted surface exposed loop (Fig. 2D). We found that TcdC(C51S/S175C)-3xmyc was efficiently biotinylated in the presence of MPB, despite lower expression than the TcdC-3xmyc (Fig 2C), showing the extracellular localization of the TcdC OB-fold.

Together, the SCAM experiments suggest that C51 and the OB-fold (C175) are located extracellularly. We sought to confirm these results in an independent assay.

### HiBiT^opt^ assay for *C. difficile* confirms extracellular location of the TcdC C-terminal domain

Previously, we have successfully used luciferase reporter assays to assess promoter activity and *in vivo* protein-protein interactions in *C. difficile* (29, 30). Here, we extend the luciferase toolbox for *C. difficile* by validating an adaptation of the Nano-Glo^®^ HiBiT Extracellular Detection System (Promega) (31).

Similar to the SmBit, the HiBiT tag is a small 11 amino acid peptide that binds to larger subunit called LgBiT to reconstitute a functional luciferase (30-32). However, in contrast to the SmBit, HiBiT has been engineered for high affinity for the LgBiT subunit (32). Due to the molecular weight (19 kDa), LgBiT extracellularly added cannot enter the cell. Thus, a luminescent signal in the presence of the substrate furimazine is only observed if the HiBiT subunit is accessible from the extracellular environment (31).

To apply this system for detection of *C. difficile* protein topology we constructed several controls carrying codon-optimized C-terminal HiBiT (HiBiT^opt^) tags. As positive control for the detection of extracellular proteins, the Sortase B (SrtB) protein was selected (SrtB-HiBiT^opt^). Sortases are membrane-anchored enzymes which catalyze the cleavage and transpeptidation of specific substrates and therefore facilitate their attachment within the cell wall (33). The genome of *C. difficile* strain 630 (and also its derivative 630Δ*erm*) has a single sortase, SrtB, present at the *C. difficile* cell wall (34, 35). The localization of SrtB and its substrates at the *C. difficile* cell surface makes SrtB a suitable candidate for the extracellular detection of the reconstituted luminescent signal. As negative control the HupA protein was used (HupA-HiBiT^opt^). This protein is a cytosolic DNA binding protein that is not secreted to the extracellular environment and thus should not be accessible to the LgBiT subunit (30). All the constructs were placed in a modular vector under the control of the anhydrotetracycline (ATc) -inducible promoter P_tet_ (28). As observed in other bioluminescence assays (29, 30) a background signal is detected from non-induced cells (Fig. 3A, T0), which is comparable to that of a medium only control (17872.8 ± 4397.7 RLU/OD; data not shown). As expected, expression of the positive control SrtB-HiBiT^opt^ leads to a 2-log increase of the luminescence signal after 45 minutes of induction (3.2×10^6^ ± 2.5×10^5^ RLU/OD, T1, Fig. 3A). No significant increase of the luminescent signal was detected in the cells expressing the negative control HupA-HiBiT^opt^, confirming that LgBiT does not enter *C. difficile* cells. The lack of signal is not due poor induction, as a clear signal is visible in the lysates of the induced samples for both the SrtB-HiBiT^opt^ and HupA-HiBiT^opt^, at the expected molecular weights of 26 kDa and 12kDa, respectively (Fig. 3B). We conclude that the HiBiT^opt^ system as employed is suitable for determining the subcellular localization of the C-terminal domain of *C. difficile* proteins.

**Figure 3.**
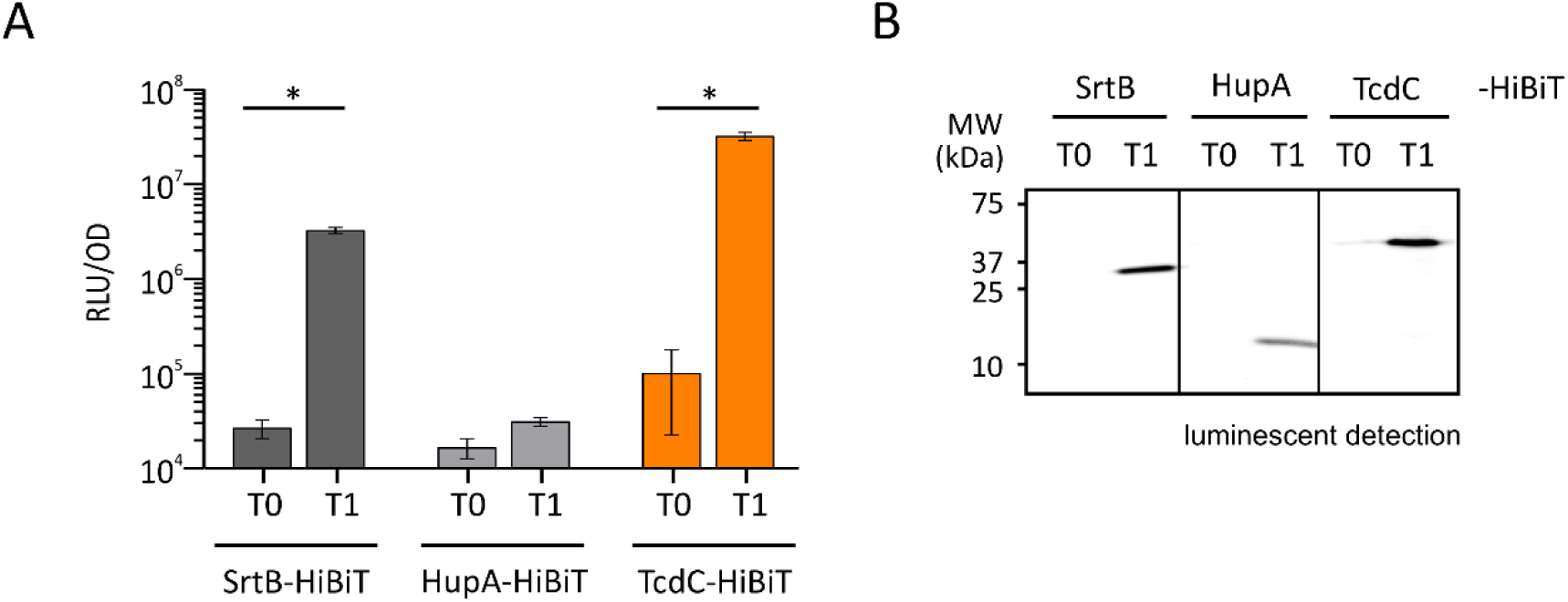
Detection of C-terminal HiBiT^opt^ tags. **A)** Proteins of interest were C-terminally fused to a HiBiT protein tag and induced with 50 ng/mL ATc for 45 minutes. Optical density-normalized luciferase activity (RLU/OD) is shown right before induction (T0) and after 45 min of induction (T1). HiBiT^opt^-tagged sortase (dark grey bars) and HupA (light grey bars) proteins were used as extracellular and intracellular controls, respectively. TcdC-HiBiT^opt^ associated luciferase activity is displayed in the orange bars. The averages of biological quadruplicate measurements are shown, with error bars indicating the standard deviation from the mean. Significance was defined as higher than p<0.001(*) by two-way ANOVA. **B)** Blot detection of HiBiT^opt^- tagged proteins resolved on a 12% SDS-PAGE. Sample volumes were normalized for optical density of the cultures from which they were derived. Expression of HiBiT^opt^ fused proteins was observed at 0 (T0) and 45 minutes after induction (T1).

Next, HiBiT^opt^ was fused to the C-terminus of TcdC (TcdC-HiBiT^opt^). We observed a 3-log increase in the luminescent signal after 45 min of induction (3.2×10^7^ ± 3.0×10^6^ RLU/OD at T1, Fig. 3A). Expression of the TcdC-HiBiT^opt^ was confirmed by detection of a clear signal at the expected MW of approximately 39 kDa (Fig. 3B). We observe a low level signal for the non-induced TcdC-HiBiT^opt^ expression construct, both in the luciferase assay (T0, Fig. 3A)_and in the detection of tagged protein (T0, Fig. 3B), which was not observed for SrtB or HupA. As all proteins are expressed from the same promoter, this possibly indicates more efficient translation, and thus higher expression, of TcdC-HiBiT^opt^ under non-inducing conditions. Alternatively, differences in luciferase detection levels might be explained by the accessibility of the HiBiT^opt^ fusion proteins for the LgBiT subunit, which in turn is affected by the structure and the exact localization of the proteins.

Taken together, the HiBiT^opt^ experiments indicate that the C-terminus of TcdC is located in the extracellular environment (like SrtB) and not in the intracellular environment (like HupA).

## Discussion

The importance of TcdC for regulation of toxin expression is highly controversial. Though the prevailing model suggests that TcdC is an anti-sigma factor with a role as a negative regulator of toxin transcription (19, 20), several other studies have found no clear relation between TcdC expression and the toxin levels (17, 18, 21, 22). Previous biochemical analyses of TcdC revealed that it is membrane-associated and that the C-terminus of TcdC comprises a dimerization domain and a domain with a predicted OB-fold, that is important for transcriptional repression (10). However, the localization of the C-terminus of TcdC has not been studied to date and this was addressed in the present study. Three independent lines of evidence suggest an extracellular location for the OB-fold domain of TcdC.

*In silico* analyses of the TcdC amino acid sequence using TMHMM 2.0 (24), TOPCONS (25), and SignalP 5.0 (26) suggest the presence of a transmembrane helix, but predicts no high-probability cleavage as expected for a secretion signal for any of the canonical secretion pathways (Fig. 1). The prediction of a transmembrane domain is consistent with the previously described biochemical assays demonstrating association of TcdC with the *C. difficile* membrane (16, 23). Analysis did not reach consensus on the localization of the N-terminus, due to low reliability scores and differences obtained with the prediction methods (Fig. 1). Nevertheless, both TMHMM 2.0 and TOPCONS suggest that the C-terminus of TcdC (residues 50 to 232, Fig. 1) is located on the outside of the cell. The *in silico* predictions are supported by the results of the Scanning Cysteine Accessibility Method (SCAM) and the HiBiT^opt^ experiments.

For the SCAM experiments we took advantage of the 2 cysteine residues that are natively present after the transmembrane helix of TcdC. Biotin labelling of TcdC-3xmyc was detected, suggesting one or both of the native cysteine residues are accessible for labelling by MPB (27) and located in the extracellular environment (Fig. 2C). TcdC(C51S)-3xmyc did not show any MBP labelling (Fig. 2C) although it contains the cysteine at position 184, in the 4^th^ ß-sheet of the OB-fold domain. The compact fold of this domain might interfere with MPB labelling, though we cannot exclude other reasons. As TcdC-3xmyc labelling was solely due to the presence of the native cysteine 51, downstream of the predicted transmembrane domain (Fig. 1 and 2B) and we were interested in validating the location of the OB-fold domain, we used the sequence of TcdC(C51S)-3xmyc as a template to introduce a cysteine residue at position 175. The native serine at position 175 is located in a loop connecting ß-sheet 3 and 4 in the OB-fold and predicted to be solvent exposed, making this position an ideal candidate for MPB labeling (Fig. 2). When expressing TcdC(C51S/S175C)-3xmyc, efficient labeling was observed, indicating that lack of labelling is not a generic feature of the OB-fold and supporting the extracellular location of the TcdC C-terminus (Fig. 2C). We note that the pattern of labelling is different between the TcdC-3xmyc and TcdC(C51S/S175C)-3xmyc. We do not know the nature of these differences, but as all bands appear to run at a higher molecular weight than expected from the theoretical mass of TcdC (as also observed by others (10, 23)), they may represent post-translational modifications. Glycosylation is one of the most common post-translational modifications found in several bacteria, particularly at the cell surface (36). Along with the TcdC extracellular localization, the higher molecular forms observed could be glycosylation forms of the protein. Our experiments do not allow us to determine the possible effects of replacing the serine and cysteine residues on the structural stability of TcdC and/or its interactions with binding partners (37). Nevertheless, our results are confirmed in independent experiments using the HiBiT^opt^ system, which to our knowledge, is applied for the first time in *C. difficile* here and extends our existing luciferase toolkit (29, 30).

Our finding that the TcdC C-terminus is extracellular challenges the prevailing model of TcdC as an anti-sigma factor. Anti-sigma factors generally sequester their cognate sigma factors away from RNAP using substantial cytoplasmic domains (38). The small N-terminal sequence, that may or may not be intracellular, is not likely to fulfill this function. Our experiments clearly place the C-terminal domain, that previously was postulated to be responsible for transcriptional repression (10), in a different environment than TcdR and RNAP. One has to wonder whether the OB-fold would ever be in contact with these proteins, as was demonstrated in experiments using purified proteins (outside the context of a *C. difficile* cell) and in heterologous expression systems (10). It should be noted that an extracellular location of TcdC does not exclude a function as a negative regulator of toxin production, but if so, suggests that it does so through an indirect mechanism.

Our data rather implies binding of OB-fold to an extracellular ligand. Bacterial OB-fold proteins have been identified in bacterial genomes and can bind a wide variety of molecules (39, 40). Thus, TcdC might bind extracellular oligonucleotides and/or oligosaccharides. It has been previously shown that the OB-fold is able to bind G-quadruplex structures, but the relevance of this binding has yet to be determined and it is conceivable that G-quadruplex binding mimics binding of its natural substrate as proposed earlier (23). In the extracellular environment, TcdC might bind oligosaccharides or extracellular DNA (eDNA) (41-43). In *Staphylococcus aureus* the sAg-like protein 10 (SSL10) binds to human IgG1 Fc primarily by its N-terminal OB-fold domain and can play a role during *S. aureus* infections (44). In *Salmonella typhimurium,* the small periplasmic protein YdeI contains an OB-fold domain and contributes to the resistance to antimicrobial peptides by interaction with the OmpF porin (45). The VisP protein, a protein from the bacterial oligonucleotide/oligosaccharide-binding fold family also present in S. *typhimurium*, binds to the peptidoglycan sugars and also to the inner membrane protein LpxO, mediating resistance and pathogenesis in *S. typhimurium* (46). To identify the TcdC ligand(s), a cross-linking and subsequent mass-spectrometry based method with tagged TcdC could be used. In addition, structural studies of TcdC and its ligands could show how the OB fold of TcdC has evolved and to what extend TcdC contributes to downregulation of the large *C. difficile* toxins.

In summary, this study indicates an extracellular localization of the C-terminus of TcdC, which is incompatible with its proposed function as an anti-sigma factor. Further studies are required to elucidate the role of TcdC in *C. difficile* development and toxin regulation.

## Materials and Methods

### Topology prediction

To determine the topology of *C. difficile* TcdC protein (UniProt ID: Q189K7) the amino acid sequence was analyzed by two computer programs for transmembrane and topology assessment: TMHMM 2.0 (http://www.cbs.dtu.dk/services/TMHMM-2.0) (24), with extensive output format, with graphics; and TOPCON 2.0 (http://topcons.cbr.su.se/) (25). The TcdC sequence was analyzed with SignalP 5.0 program, for signal peptide prediction, with long output for gram-positive organisms, (http://www.cbs.dtu.dk/services/SignalP/)(26). All the results were visualized with GraphPad Prism 8 software (version 8.1.2)

### Strains and growth conditions

*E. coli* strains were grown aerobically at 37°C in Luria Bertani broth (LB, Affymetrix) supplemented with chloramphenicol at 15 µg/mL or 50 µg/mL kanamycin when required. Plasmids (Table 1) were maintained in *E. coli* strain DH5α and transformed using standard procedures (47). *E. coli* CA434 (48), was used for plasmid conjugation with *C. difficile* strain 630Δ*erm* (34, 49).

**Table 1.** Plasmids used in this study.

| Name | Relevant features * | Source/Reference |
| --- | --- | --- |
| pCR2.1-TOPO | TA vector; pMB1 oriR; *km amp* | ThermoFisher |
| pCR2.1TcdC | pCR2.1-TOPO with *tcdC*; *km amp* | This study |
| pRPF185 | *tetR* P<sub>tet</sub>-*gusA*; *catP* | (28) |
| pLDJ1 | *tetR* P<sub>tet</sub>-*tcdC*-3xmyc; *catP* | This study |
| pLDJ2 | *tetR* P<sub>tet</sub>-*tcdC*(C51S)-3xmyc; *catP* | This study |
| pLDJ5 | *tetR* P<sub>tet</sub>-*tcdC*(C51S/S175C)-3xmyc; *catP* | This study |
| pJC111 | *tetR* P<sub>tet</sub>-*tcdC*-*hiBiT*<sup>opt</sup>; *catP* | This study |
| pAF302 | *tetR* P<sub>tet</sub>-*hupA*-*hiBiT*<sup>opt</sup>; *catP* | This study |
| pAP233 | *tetR* P<sub>tet</sub>-*srtB*-*hiBiT*<sup>opt</sup>; *catP* | This study |

\* *amp* – ampicillin resistance cassette, *km* – kanamycin resistance cassette, *catP* – chloramphenicol resistance cassette

*C. difficile* strains grown anaerobically in Brain Heart Infusion broth (BHI, Oxoid), with 0,5 % w/v yeast extract (Sigma-Aldrich), supplemented with *Clostridium difficile* Selective Supplement (CDSS; Oxoid) and 15 µg/mL thiamphenicol when necessary, at 37°C in a Don Whitley VA-1000 workstation or a Baker Ruskin Concept 1000 workstation with an atmosphere of 10% H_2_, 10% CO_2_ and 80% N_2_.

The growth was monitored by optical density reading at 600 nm (OD_600_).

All strains are described in Table 2.

**Table 2.** *C. difficile* strains used in this study.

| Name | Relevant Genotype/Phenotype* | Origin /Reference |
| --- | --- | --- |
| 630Δ*erm* | *C. difficile* 630Δ*erm*; Erm<sup>S</sup> | (34, 49) |
| LDJ1 | 630Δ*erm* pLDJ1; Thia<sup>R</sup> | This study |
| LDJ2 | 630Δ*erm* pLDJ2; Thia<sup>R</sup> | This study |
| LDJ5 | 630Δ*erm* pLDJ5; Thia<sup>R</sup> | This study |
| JC267 | 630Δ*erm* pJC111; Thia<sup>R</sup> | This study |
| JC271 | 630Δ*erm* pAF302; Thia<sup>R</sup> | This study |
| AP239 | 630Δ*erm* pAP233; Thia<sup>R</sup> | This study |

\* Erm<sup>S</sup> – Erythromycin sensitive, Thia<sup>R</sup> – Thiamphenicol resistant

### Strain construction

All oligonucleotides from this study are listed in Table 3. All PCRs were performed on 630Δ*erm* genomic DNA (34), unless indicated otherwise. Expression vectors are all based on pRPF185 (28). All DNA sequences in the recombinant plasmids were verified by Sanger sequencing of the region of the plasmid encompassing the inserted fragment and the full anhydrotetracycline-inducible promoter.

**Table 3.**
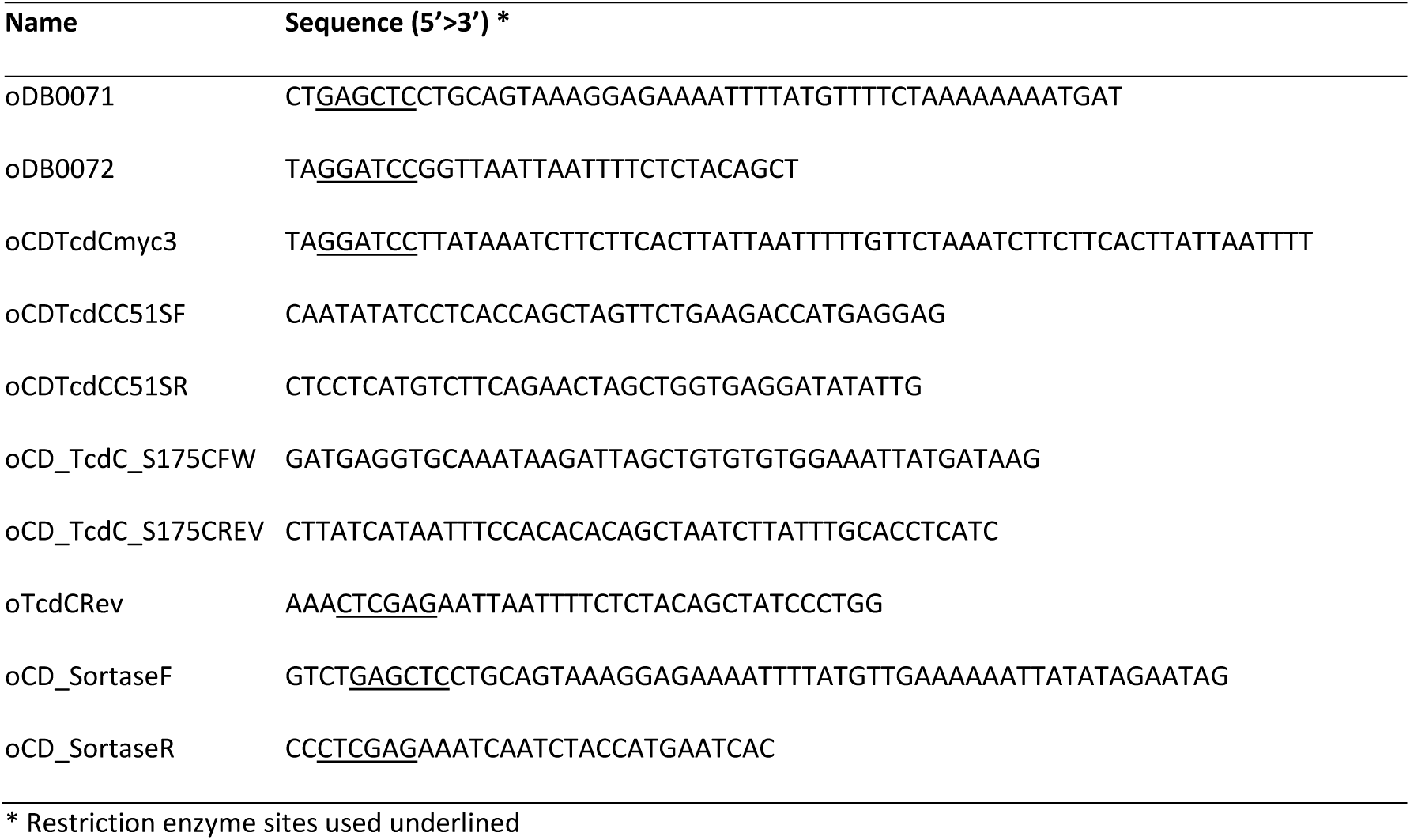
Oligonucleotides used in this study.

#### Construction of the *tcdC* mutants for SCAM analysis

To construct the expression constructs for the *tcdC* mutants, the *tcdC* gene (CD0664 from *C. difficile* 630; GenBank accession no. YP_001087138.1) was amplified by PCR using primers oDB0071 and oDB0072 from *C. difficile* chromosomal DNA. The PCR product was subsequently cloned into pCR2.1TOPO (Invitrogen), according to manufacturer’s instructions, to yield vector pCR2.1TcdC (Table 1). The TcdC fragment was amplified from pCR2.1TcdC with primers oDB0071 and oCDTcdCmyc3, which allows the addition of a 3xmyc-tag at the C-terminus. The resulting PCR fragment was digested with SacI and BamHI and ligated into pRPF185 digested with the same enzymes, to yield vector pLDJ1 (Table 2), placing *tcdC* under the control of the anhydrotetracycline inducible promoter (P_tet_).

Site-specific mutations were introduced in pCR2.1TcdC sequence via QuikChange mutagenesis (Stratagene). For the C51S mutation, primer set oCDTcdCC51SF/oCDTcdCC51SR was used. Addition of a Myc-tag and cloning into pRPF185 was performed as described for the wild type protein, yielding pLDJ2 (Table 1).

Vector pLDJ2, harbouring P_tet_-*tcdC(C51S)-3xmyc* was used as template to generate the mutant *tcdC*(C51S/S175C) via QuikChange mutagenesis (Stratagene) using the primer set oCDTcdCS175CFW/oCDTcdCS175CREV, yielding pLDJ5 (Table 1).

#### Construction of the HiBiT^opt^ fusions

The *hupA* gene (CD3496 from *C. difficile* 630; GenBank accession no. NC_009089.1) fused at the 3’ end to the *hibiT*^*opt*^ codon optimized coding sequence (HupA-HiBiT^opt^) was synthesized by Integrated DNA Technologies, Inc. (IDT). The synthesized fragment (full sequence available as Supplementary Data) was digested with BamHI and SacI, and cloned into pRPF185 digested with same enzymes, yielding vector pAF302 (Table 1).

The *tcdC* gene was amplified by PCR with primer set oDB0071/oTcdCRev (Table 3) from *C. difficile* genomic DNA). The PCR fragment was digested with XhoI and SacI and cloned into similarly digested pAF302, yielding vector pJC111 (Table 1).

The *srtB gene* (CD2718 from *C. difficile* 630; GenBank accession no. YP_001089230.1) was amplified from *C. difficile* genomic DNA with primers oCD-sortaseF/oCD-sortaseR (Table 3), digested with XhoI and SacI, and placed into similarly digested pAF302, yielding vector pAP233 (Table 1).

### Affinity purification of the anti-TcdC polyclonal antibodies

Polyclonal antibodies against TcdC were raised in rabbits using the peptide CQLARTPDDYKYKKV (17). To reduce background signals, serum from the final bleed from the immunized rabbits was subjected to affinity purification. Recombinant soluble 10xHis-TcdCΔN50 (lacking the N-terminal 50 amino acids) (23) was blotted on a PVDF membrane. The blot was stained with Ponceau S solution (0,2% (w/v) Ponceau S, 1% acetic acid) for 5 minutes. Subsequently, the blot was washed with TBST (500 mM NaCl, 20 mM Tris, pH 7,4, 0,05% (v/v) Tween-20) until the TcdC band was visible. The band was cut out of the blot and the piece of membrane was washed with Ponceau destaining solution (PBS, 0,1% NaOH) for 1 minute. Subsequently, the piece of blot was washed twice with TBST for 5 minutes. Then, the membrane was soaked in acidic glycine buffer (100 mM glycine, pH 2,5) for 5 minutes and washed twice in TBST for 5 minutes. The blot was blocked in 10% ELK (in TBST) for 1 hour at room temperature. After washing it twice with TBST for 5 minutes, the blot was incubated with 10 ml of 5 x diluted serum (in TBS) overnight at 4 °C. Afterwards, the diluted serum was removed and the blot was washed 3 times with TBST for 5 minutes and 2 times with PBS for 5 minutes. The blot was incubated with 1 mL of acidic glycine buffer for 10 minutes to elute the antibodies. The eluted antibody solution was immediately neutralized by adding 1M Tris-HCl, pH 8.0. After addition of sodium azide (5 mM) and BSA (1 mg/ml), the affinity purified antibodies were stored at 4°C until use in experiments.

### Substituted-Cysteine Accessibility method

To perform the Substituted-Cysteine Accessibility Method (SCAM), 25 ml *C. difficile* cultures were induced with 200 ng/ml anhydrotetracycline (ATc) at an OD_600_ of 0.3, for 2 hours. The cultures were centrifuged for 10 min at 2800 × g and the pellet was frozen at -20°C when needed. Subsequently, the pellet was resuspended in 600 μl GTE buffer (50 mM glucose; 20 mM tris-HCl; 1 mM EDTA, pH7.5) supplemented with Complete protease inhibitor cocktail (CPIC, Roche Applied Science) and aliquoted in 100 μl. For the cysteine labelling, 1 mM of MPB was added and the samples were incubated for 15 min on ice. The reactions were quenched by the addition of 73 mM β-mercaptoethanol. Samples were washed twice in GTE buffer + CPIC and centrifuged at 2800 x g. Pellets were resuspended in 100 μl solubilization buffer (50 mm Tris-HCl, pH 8.1, 2% SDS, 1mM EDTA) with mixing for 5 min and sonication (2 × 5-10 sec pulses). To remove unspecific biding, 400 μl 0.2% PBS-T + Triton X-100 + CIPC was added to the sample together with 30 μl 50% protein A sepharose CL-4B (Amersham) slurry in phosphate buffered saline (PBS) supplemented with 1% bovine serum albumin (BSA), previously equilibrated in PBS/BSA 1%. After overnight incubation at 4°C, Protein A sepharose beads were removed by gentle centrifugation.

For immunoprecipitation, 50 μl of 50% protein A sepharose CL-4B (Amersham) slurry was added to each sample with the affinity purified polyclonal rabbit anti-TcdC antibody (1:200) and incubated at 4°C with gentle mixing for 2 hours. The slurry was pelleted (4000 × g) and washed 2 times with TENT buffer (150 mm NaCl; 5 Mm EDTA; 50 mm Tris; Triton-X-100 0.5%; pH 7.5) + 1% BSA + 0.5 M NaCl, 2 times with TENT + 1% BSA + 0.25M NaCl and 2 times with TENT buffer. The pellet was resuspended in 50 μl SDS loading buffer (250 mM Tris-Cl pH 6.8, 10 % SDS, 10% β-mercaptoethanol, 50% glycerol, 0.1% bromophenol blue) and incubated at 50°C for 5 min. Samples were spun down prior to SDS-PAGE analysis.

### Immunoblotting and detection

Proteins were separated on a 12% SDS-PAGE gel and transferred onto polyvinyl difluoride (PVDF) membranes (Amersham), according to the manufacturers’ instructions. The membranes were probed with monoclonal mouse anti-myc antibodies (α-myc, 1:1500, Invitrogen) or mouse anti-biotin- (α-biotin, 1:1000) in PBST. After washing the blots with PBST, a secondary goat-anti-mouse HRP antibody (1:1000, Dako) was used. Bands were visualized using the Clarity ECL Western Blotting Substrates (Bio-Rad) on an Alliance Q9 Advanced machine (Uvitec).

### HiBiT^opt^ Assay

*C. difficile* cells were induced with 50 ng/mL ATc at an OD_600_ of 0.3-0.4, for 45 min. An one ml sample was collected and centrifuged (4000 × g) for SDS-PAGE analysis and luminescent detection of HiBiT^opt^-tagged proteins on a blot.

To measure luciferase activity, 50 µL sample was incubated with 50 µL Nano-Glo HiBiT Extracellular Detection System, a mixture of the NanoLuc LgBiT protein and luciferase substrate in buffer (Promega) for 5 min, in a 96-well white F-bottom plate. Luciferase activity was measured on a GloMax Multi+ instrument (Promega), with a 0.5 s integration time. All luciferase measurements were taken immediately after sampling. Data was normalized to OD_600_ of the culture the samples were derived from and statistical analysis was performed with Prism 7 (GraphPad Inc, La Jolla, CA) by two-way ANOVA.

For luminescent detection of HiBiT^opt^-tagged proteins on a blot, total protein was resolved on a 12% SDS-PAGE gel and transferred onto polyvinyl difluoride (PVDF) membranes (Amersham). The membranes were washed with TBST and incubated with 200-fold diluted LgBiT protein in TBST (N2421, Promega), for 1 hour at room temperature. 500-fold diluted Nano-Glo Luciferase Assay Substrate (N2421, Promega) was added and incubated for 5 min at room temperature with gentle shaking. The membranes were analysed using an Alliance Q9 Advanced machine (Uvitec).

### Software

Structural analysis was performed using PyMOL Molecular Graphics System, Version 1.76.6 Schrödinger, LLC. GraphPad Prism 8 (version 8.1.2) was used for statistical analysis. Images were prepared for publication in CorelDRAW Graphics Suite X7 software.

## Acknowledgments

Work in the group of WKS is supported by a Vidi Fellowship (864.10.003) of the Netherlands Organization for Scientific Research (NWO) and a Gisela Their Fellowship from the Leiden University Medical Center.

## Supplementary Data

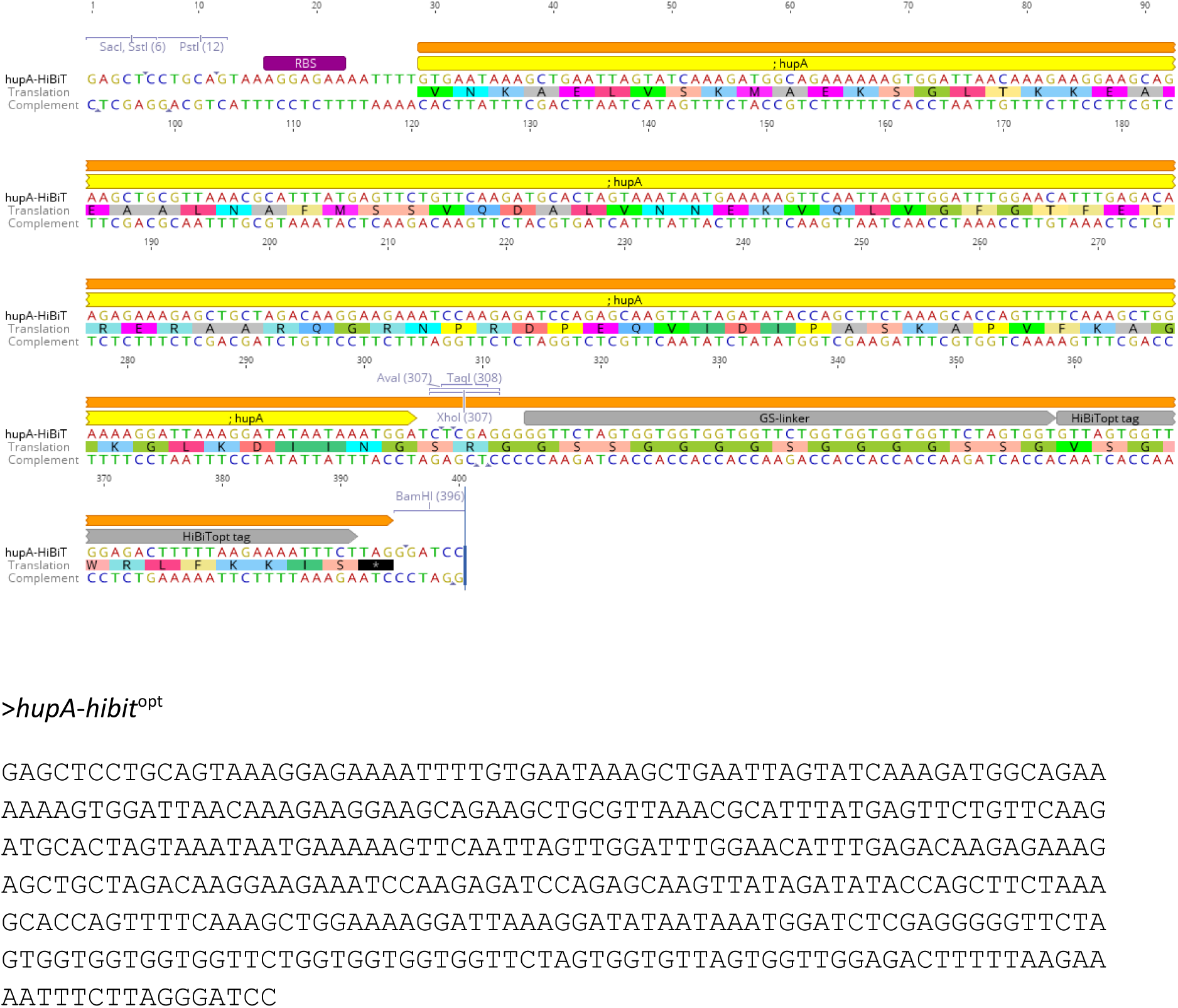

